# Two Sites, Two Languages: Dual-site tDCS and EEG Evidence for L1 Feature Transfer in L2

**DOI:** 10.1101/2025.06.03.657565

**Authors:** Alaa M. Salem, Daniel Gallagher, Emi Yamada, Shinri Ohta

## Abstract

The neural bases of syntactic and semantic processing remain unclear. While prior transcranial direct current stimulation (tDCS) studies have targeted either the inferior frontal gyrus (LIFG) or left superior temporal gyrus (LSTG), we test whether dual stimulation of both alters second language (L2) anomaly based on native language (L1). Chinese and Korean participants evaluated Japanese sentence correctness; Japanese shares morpho-syntax with Korean and semantic radicals with Chinese. For the N400, dual anodal tDCS elicited a significant interaction stimulation and L1 (*F* (1,960.7) = 5.48, *p* = 0.0194), stimulation and sentence type (*F* (5,960.7) = 2.28, *p* = .0448), and L1 and sentence type (*F* (5, 960) = 2.28, *p* = 0.045). For the P600, a significant effect of stimulation and L1 interaction (*F* (1,1303.1) = 9.86, *p* = 0.0017), and stimulation x sentence type (*F* (5,1303.1) = 2.35, *p* = 0.039), suggesting L1 typology affected semantic integration during dual-stimulation tDCS.

## Main

The brain is a profoundly complex organ, and while certain regions are known to be more active than others during specific cognitive tasks, it is a misinterpretation to name complex cognitive functions to isolated brain areas. Alternatively, these regions often cooperate with other regions or parts of the brain (Cocchi et al., 2013). Consequently, this forms sophisticated networks within the brain that support various cognitive processes. For instance, while the left inferior frontal gyrus (LIFG) (Sheng et al., 2019) and the left superior temporal gyrus (LSTG) (Berken et al., 2016) are extensively involved in language processing, they do not work in isolation from each other or any other region. The LIFG, also known as Broca’s area, plays a key role in syntactic processing and language production. On the other hand, the LSTG, often known as Wernicke’s area, is crucial for semantic processing and language comprehension by deciphering the linguistic inputs. Nevertheless, these regions collaborate with other brain parts, such as the prefrontal cortex, parietal lobes, and subcortical structures, to create a comprehensive language processing network (Friederici, 2011).

Brain stimulation techniques such as transcranial magnetic stimulation and transcranial direct/alternating current stimulation (tDCS/tACS) can function as inhibitors or activators of brain activity using magnetic or electrical methods. They can be applied in various ways, such as treating neurological or psychiatric diseases, addressing health-related issues, and advancing neuroscience research to learn more about the brain (Paulus, 2014). Advances in non-invasive brain stimulation techniques, such as tDCS and the more advanced high-definition tDCS, have opened new avenues for exploring various aspects of language learning, its complex processing, and other cognitive functions. Anodal tDCS, which modulates cortical excitability by increasing neuronal firing rates (Das et al., 2016), has contributed significantly to our understanding of the functional specialization of cortical regions. For example, stimulation of the LIFG has been shown to enhance complex sentence comprehension (Krause et al., 2023), while stimulation of the LSTG has been associated with improved learning of novel abstract and concrete lexical items (Kurmakaeva et al., 2021). Although these regions have been extensively studied in isolation, the effects of dual-site application of tDCS—particularly its effects on syntactic and semantic violations—remain underexplored.

Electroencephalography (EEG) is a non-invasive method to record the brain’s electrical activity, providing temporally accurate information about neural oscillations and brain states (Gevins et al., 1999). When combined with tDCS, EEG can help reveal the neurophysiological mechanisms underlying tDCS-induced modulation of brain activity. Studies have shown that anodal tDCS can increase cortical excitability, leading to changes in EEG power across various frequency bands, such as theta, alpha, beta, and gamma (Ke et al., 2023). This EEG-tDCS combination allows for the observation of real-time changes in brain activity, enhancing our understanding of how tDCS influences cognitive functions, particularly in language processing (Bhattacharjee et al., 2024). By applying dual-site anodal tDCS simultaneously over the LIFG and LSTG, the present study seeks to bridge the gap in understanding the interconnected roles of these regions. This dual-site approach allows for a more comprehensive understanding of how syntactic and semantic processing are modulated in second language (L2) learners.

L2 acquisition provides additional insights into neural plasticity and dynamics. Cross-linguistic influences from one’s native language (L1) significantly shape the learning phases, pronunciation, and processing of L2. These influences are particularly prominent in tasks involving syntactic and semantic anomaly detection (Hahne & Friederici, 2001). In the current study, we evaluate the semantic and syntactic naturalness of Japanese sentences among late L2 learners with varying L1 backgrounds (i.e., Korean and Chinese).

The P600 is a positive-going waveform typically seen in syntactic anomalies or complex syntactic processing, often surging after approximately 600 ms (Osterhout et al., 1994). On the other hand, the N400 is a negative-going waveform typically seen in semantic anomalies or complex semantic processing, often emerging around 400 ms post-stimulus onset (Kutas & Hillyard, 1984). Multi-stream models propose that event-related potential (ERP), N400, and P600 processing occur in parallel and independent pathways. On the other hand, Retrieval-Integration (RI) theory assumes a single-stream sequence, where lexical retrieval (N400) takes place before compositional integration (P600) as the integration effort increases in a predictable way (Brouwer et al., 2012).

Our hypothesis was that native Korean L1 speakers learning Japanese as an L2 may exhibit enhanced syntactic processing with distinct P600 modulations, as shown by accuracy, reaction time (RT), and a more pronounced P600 in the ERP data, due to shared morpho-syntactic structures. Similarly, native Chinese L1 speakers might excel in semantic anomaly detection with enhanced N400 modulations, owing to the shared Chinese characters with semantic radicals. These cross-linguistic variations highlight the importance of considering the L1 background when investigating L2 processing mechanisms. This not only enhances our understanding of the neural mechanisms in the L2 brain but also fosters the development of educational strategies targeting late language learners. Additionally, the dual-site tDCS stimulation paradigm might offer a new perspective for developing innovative therapeutic applications for language disorders (Shahbazi et al., 2024).

In this study, we examined the effects of simultaneous dual-site anodal HD-tDCS applied to the LIFG and LSTG on L2 processing. By combining HD-tDCS and EEG, we investigated L1-specific modulations in the L2 P600 and N400.

## Results

To examine the effects of stimulation, control, semantic, and syntactic violations, verb type, and L1 on processing accuracy, we conducted a four-way 2 (L1: Chinese vs. Korean) × 2 (Stimulation: Sham vs. Active) × 2 (Verb Type: Transitive vs. Intransitive) × 3 (Violation: Control / Syntactic / Semantic) mixed ANOVA. We found a significant main effect of violation type, *F* (1.51, 37.87) = 38.41, *p* < .001, η² = 0.38, with semantic violations (*M* = 0.84, *SD* = 0.11) evoking higher accuracy than syntactic violations (*M* = 0.53, *SD* = 0.21; *p* < .001) and controls (*M* = 0.73, *SD* = 0.15; *p* < .001).

Chinese participants performed better in judging the semantically violated sentences (d3) than both control (d1) and syntactically violated ones (d2) (vs. d2: *M*_diff_ = –0.4081, *t* (14) = 9.326, *p* < 0.001; vs. d1: *M*_diff_ = –0.2247, *t* (14) = 5.406, *p* = 0.0001). The control condition also significantly surpassed the syntactic condition (d2; *M*_diff_ = 0.1833, *t* (14) = 2.739, *p* = 0.016), showing overall performance as semantic > control > syntactic.

In contrast, Korean participants showed no significant difference between the control and semantically violated sentences (*M*_diff_ = –0.0006, *t* (11) = 0.017, *p* = 0.987); however, both outperformed the syntactic condition (d3 > d2: *M*_diff_ = –0.2134, *t* (11) = 4.613, *p* = 0.0022; d1 > d2: *M*_diff_ = 0.2128, *t* (11) = 3.873, *p* = 0.0026), yielding the hierarchy: semantic ≈ control > syntactic.

These results indicate a significant interaction between participant group (Chinese vs. Korean) and condition type, especially in how syntactic, semantic, and control conditions are comparatively processed. However, there was a significant transitivity effect modulated by verb type (transitive [c1] vs. intransitive [c2]), as shown by the simple effects for the interaction between violation type and verb type—particularly in the control condition (*F* (1,25) = 10.99, *p* = 0.0028, *η²* = 0.054), with transitive verbs (*M* = 0.7587) outperforming intransitive verbs (*M* = 0.6852). There was no transitivity effect in the syntactic (d2) or semantic conditions (*p* > 0.05).

Although there was no significant interaction between language group and transitivity (*p* = 0.1112), follow-up simple effects showed group differences. In the Chinese group, transitive verbs in control sentences (c1-d1: *M* = 0.6990) significantly outperformed intransitive verbs (c2-d1: *M* = 0.6020, *p* = 0.0028), but there was no transitivity effect in syntactic or semantic conditions (*p* > 0.05). On the other hand, no significant transitivity effects were shown in any condition in the Korean group (*p* = 0.5605 for d1; *p* = 0.1988 for d3).

Post-hoc comparisons showed that transitivity modulated the condition differences as follows: For transitive verbs (c1), there were larger gaps between syntactic (d2) and semantic (d3) conditions (d2–d3: *M*_diff_ = −0.2980) compared to intransitive verbs (d2–d3: *M*_diff_ = −0.3235). For intransitive verbs (c2), the control condition (d1) showed lower accuracy than the semantic condition (d3) (d1–d3: *M*_diff_ = −0.1540, p = 0.0001). However, this difference was significantly reduced for transitive verbs (d1–d3: *M*_diff_ = −0.0713, *p* = 0.0256).

Similar to accuracy, to examine the effects of control, semantic, and syntactic violations on reaction time (RT), we used a four-way mixed-design ANOVA. First, there was a significant main effect of violation type, *F* (2, 50) = 17.17, *p* < .001, *η²* = 0.063, which revealed that syntactic processing (d2: *M* = 1.1447) induced the slowest RTs, followed by control (d1: *M* = 1.0282) and semantic (d3: *M* = 0.8934) conditions, with statistical significance observed in all pairwise comparisons (*p* < 0.05). A significant Condition × Transitivity interaction, *F* (2, 50) = 7.86, *p* = 0.001, *η²* = 0.009, revealed that the effect of sentence condition on RTs differed by verb transitivity. For intransitive verbs (c2), RTs followed the hierarchy: syntactic (d2: *M* = 1.1923) > control (d1: *M* = 1.0879) > semantic (d3: *M* = 0.8631), with statistical significance observed in all pairwise comparisons (p_adj < 0.001). In contrast, for transitive verbs (c1), RTs for syntactically violated sentences (d2: *M* = 1.1005) were longer than control (d1: *M* = 0.9770, *p* = 0.022) and semantic (d3: *M* = 0.9180, *p* = 0.002), but control and semantic conditions did not differ (*p* = 0.246).

Chinese participants (a1) showed significant transitivity effects, with slower RTs for intransitive verbs (c2) compared to transitive verbs (c1) in both control (c2–d1: *M* = 1.1007 vs. c1–d1: *M* = 1.0051; *F* (1, 14) = 10.92, *p* = 0.005, *η²* = 0.098) and syntactic conditions (c2–d2: M = 1.1868 vs. c1–d2: *M* = 1.0805; *F* (1, 14) = 8.65, *p* = 0.007, *η²* = 0.009). In contrast, Korean participants (a2) exhibited no transitivity effects in any violation condition (control: c1–d1: *M* = 0.9002 vs. c2–d1: *M* = 0.9741; *F* (1, 11) = 0.36, *p* = 0.561), with RTs remaining comparable across transitive and intransitive verbs. A marginal L1 × Transitivity interaction, *F* (1, 25) = 2.97, *p* = 0.097, *η*² = 0.001, further highlighted the cross-group difference.

## EEG Results

### Korean Sham Group: Syntactically Violated – Control

The topographic figure represents the difference wave calculated by subtracting the ERPs elicited by the control condition from those elicited by the syntactic condition in the sham Korean participant group (syntactic – control). By subtracting the control ERPs, the neural activity specifically related to syntactic processing is isolated. The scalp distribution of the difference in the ERPs between the syntactic and control conditions for Korean participants under sham stimulation is shown in Fig. 3A. An N400-compatible negativity was evident in the centro-parietal region (350–449 ms, continuing to ∼950 ms). A subsequent positivity emerged in fronto-temporal areas between ∼450 ms and 950 ms.

**Fig. 1.**
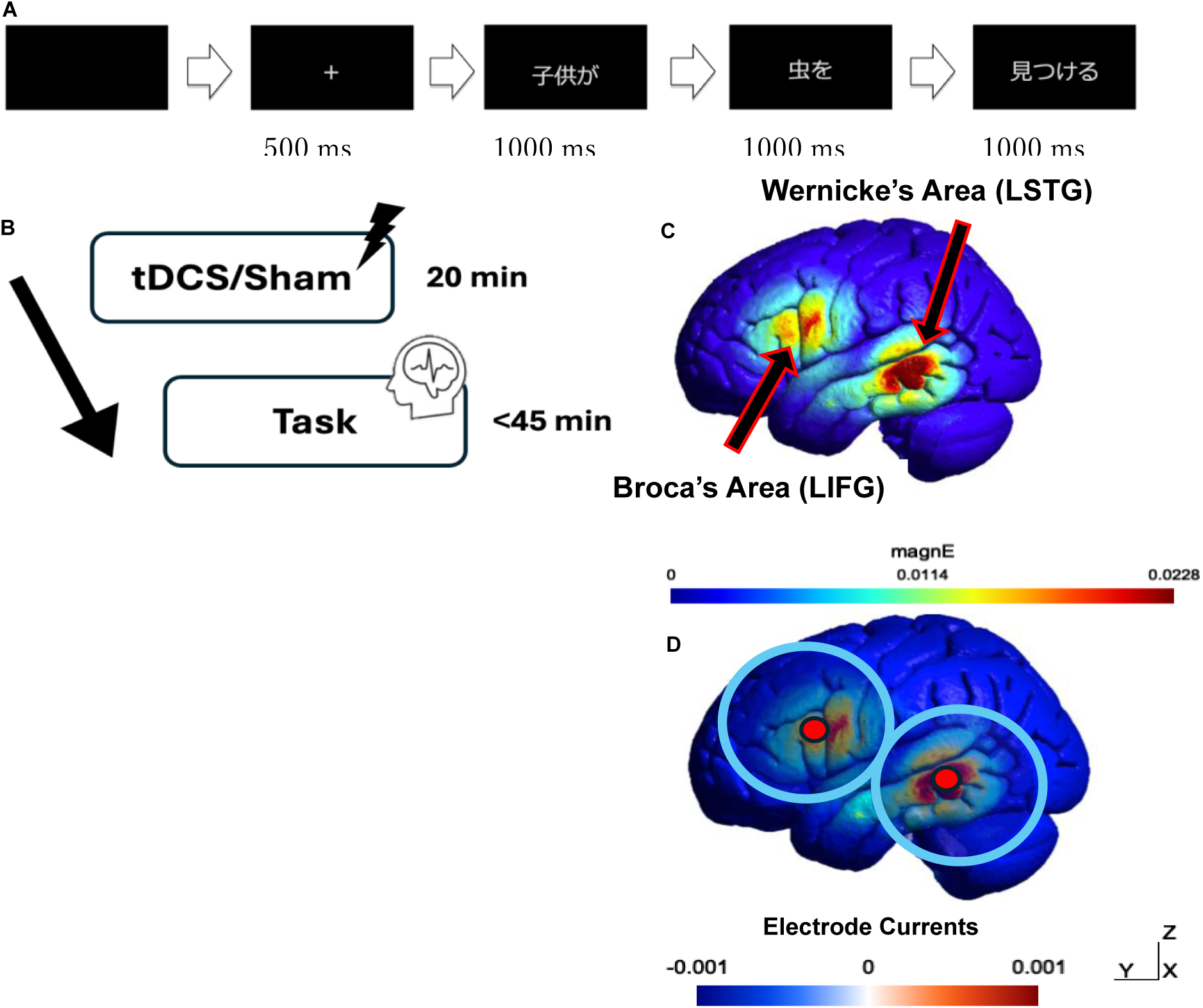
Experimental Design and Dual Anodal Simulation. A, Stimulus Presentation. **B**, Experimental Design. **C**, Dual Anodal Simulation of electric current on the canonical brain (LIFG & LSTG. **D**, Custom HD-tDCS Electrode Montage: Two large rubber-ring electrodes (blue) over the LIFG (FC5) and LSTG (TP7) served as cathodes, while two small Ag/AgCl electrodes (red) centered within each rubber-ring electrode served as anodes. SimNIBS (version 4.0.1, MNI152 space) was used for the simulation.

**Fig. 2:**
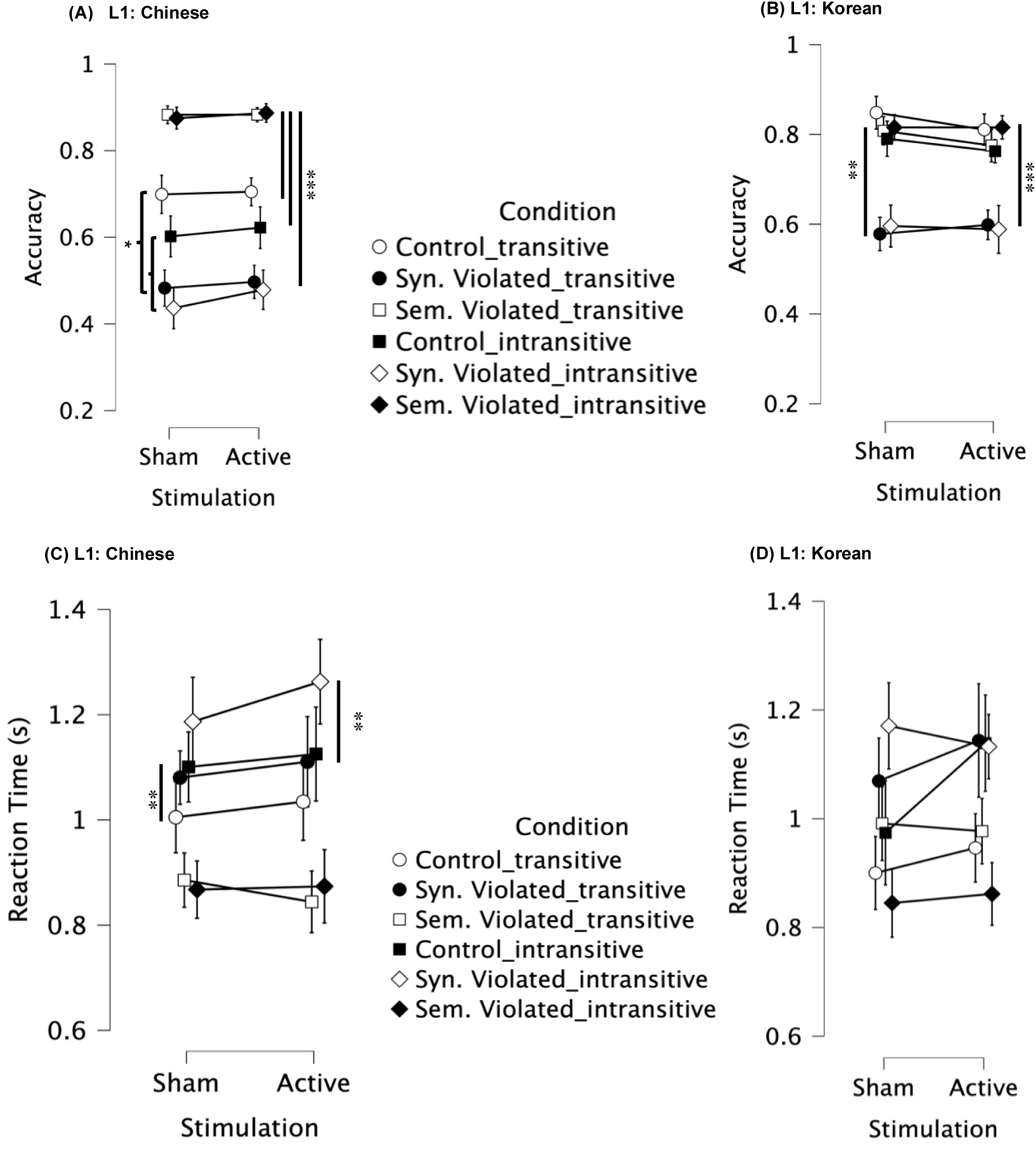
Mean Accuracy and RT by Condition, Stimulation, and Language. **A,** Mean accuracy of Chinese participants. **B**, Mean accuracy of Korean participants. **C**, Mean RT of Chinese participants. **D**, Mean RT of Korean participants.

**Fig. 3:**
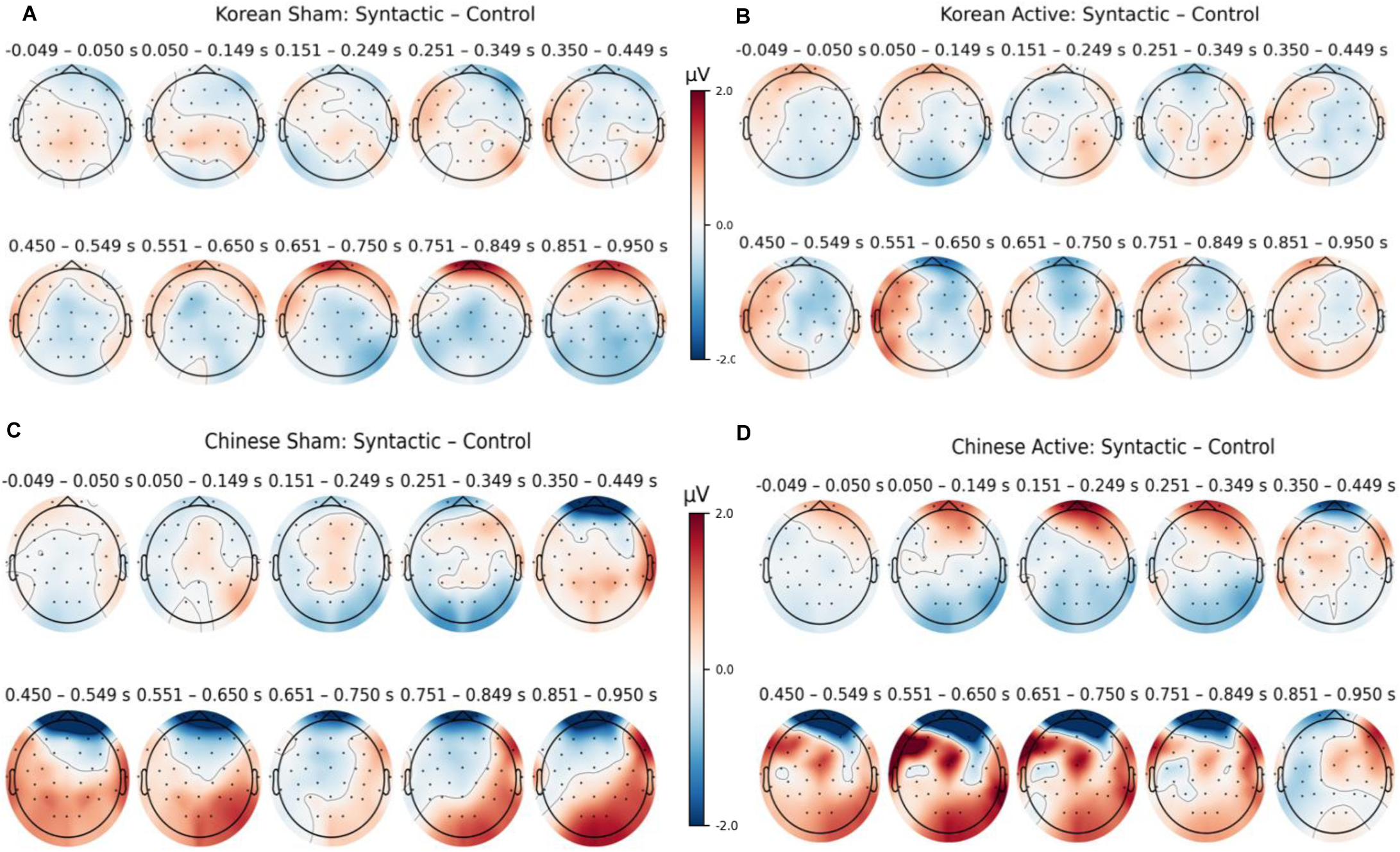
Topographical maps of the difference between syntactic and control conditions in Korean and Chinese participants. Sham (A, C) and active (B, D) stimulations. The color scale indicates the voltage difference in microvolts (μV), with red representing positive differences (relative positivity in the syntactic condition) and blue representing negative differences (relative positivity in the control condition). Each head plot represents a specific time window, as indicated above each plot. Black dots represent electrode locations.

### Korean Active Group: Syntactically Violated – Control

Similar to the sham group, the N400 was shown in the centro-parietal region, peaking around 350–449 ms. Differently, a strong positivity is observed around 450–650 ms in the left frontal and temporal region and extended to the centro-fronto-parietal regions between approximately 651 and 950 ms, indicating a persistent P600 in the active group that was not shown in the sham group (Fig. 3B).

### Chinese Sham Group: Syntactically Violated – Control

The topographic map in Fig. 3C shows positivity around 350 ms and peaking around 650 ms, as well as a negativity in the frontal region that extended to the central and left temporal region.

### Chinese Active Group: Syntactically Violated – Control

There is a noticeable frontal negativity that significantly decreased from the sham group between 350 and 449 ms. However, the positivity appeared to be more extended and stronger at the particular time 450–849 ms in the active group (Fig. 3D).

### Korean Sham Group: Semantically Violated **–** Control

In topographic Fig. 4A, the difference wave is calculated by subtracting the ERPs elicited by the control condition from those derived from the semantic condition in sham Korean participants (semantic – control). An N400 is suggested in the centro-parietal and occipital region from 350 ms and extending to right and left parieto-temporal region from 651-950ms. A positivity in the frontal and right fronto-temporal region was also shown from 450-950 ms.

**Fig. 4:**
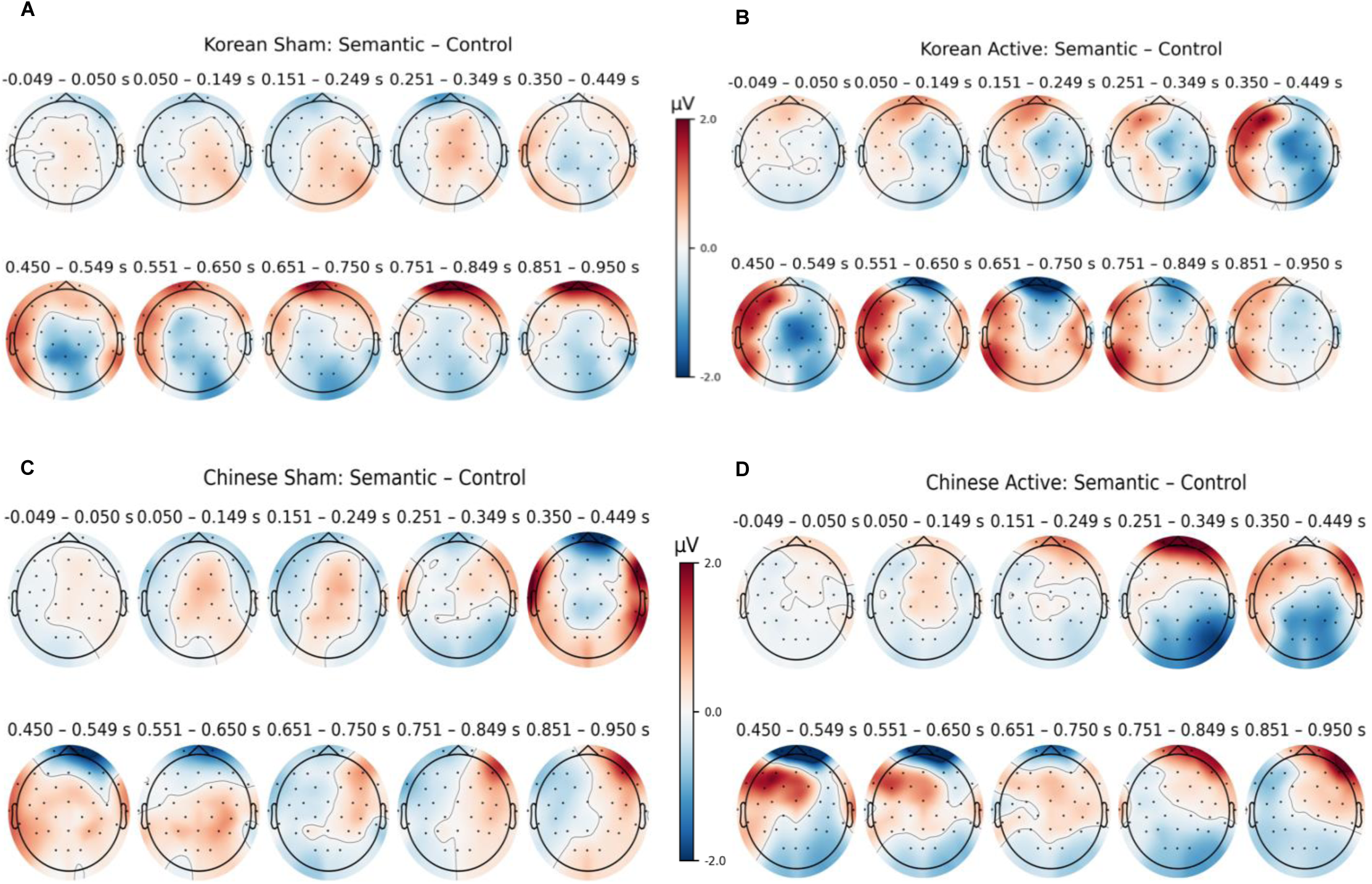
Topographical maps of the difference between semantic and control conditions in Korean and Chinese participants. Sham (A, C) and active (B, D) stimulations. The color scale indicates the voltage difference in microvolts (μV), with red representing positive differences (relative positivity in the syntactic condition) and blue representing negative differences (relative positivity in the control condition). Each head plot represents a specific time window, as indicated above each plot. Black dots represent electrode locations.

**Fig. 5.**
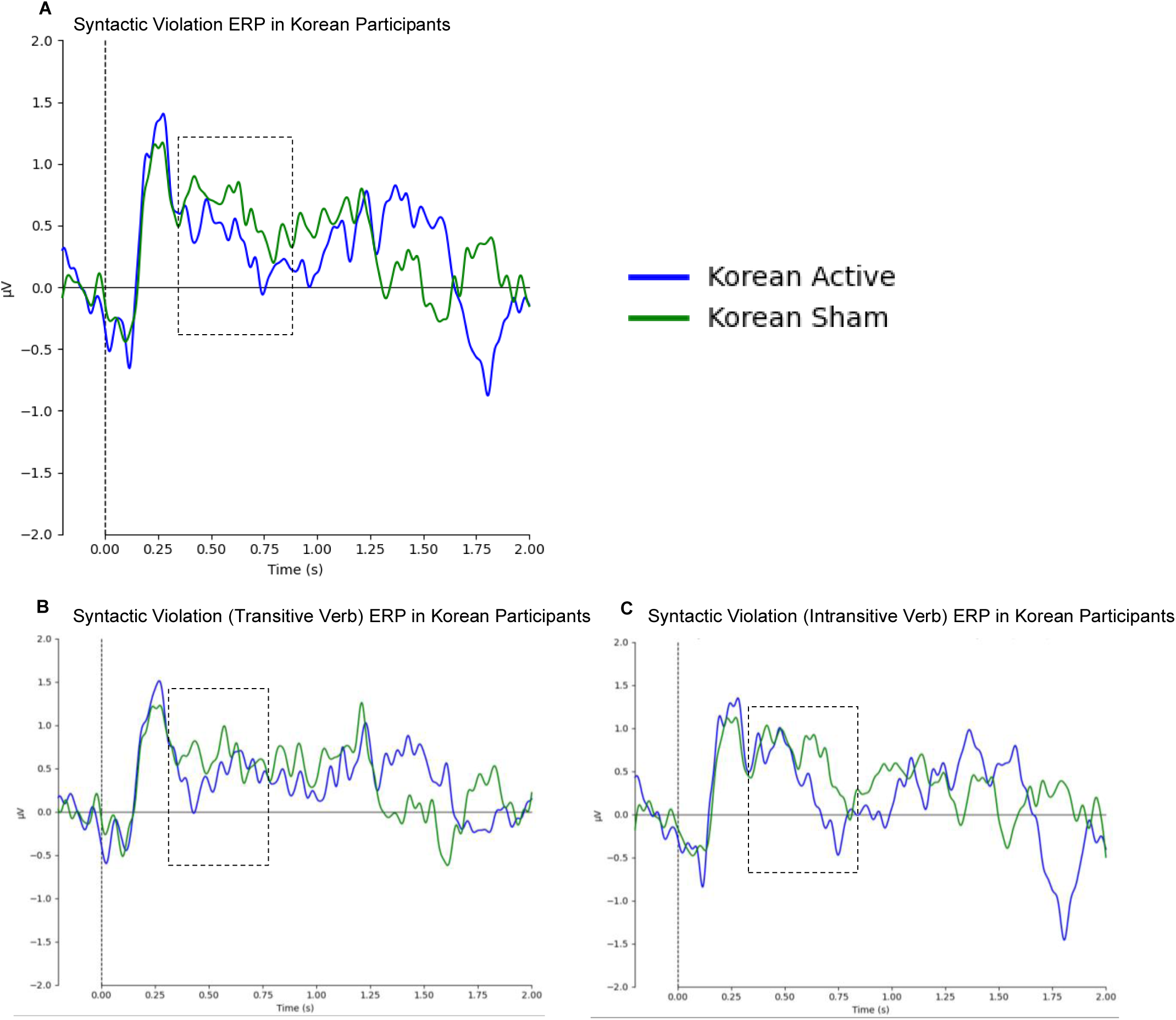
Mean grand average ERPs in Centro-parietal region in syntactic violated sentences in Korean participants (n = 12). **A**, Sham stimulation (green) shows a positivity (P600) around 500–700 ms, while under active stimulation, an N400-like waveform is observed around 300–500 ms and the P600 is significantly reduced during processing of syntactically violated sentences. **B**, Sightly reduced P600 during processing of syntactically violated sentences (transitive verbs). This might indicate that processing syntactic violations in transitive verbs is more demanding, whereas active stimulation may increase reliance on semantic information during sentence reanalysis. **C**, Active stimulation significantly reduces the P600 compared to the sham group (intransitive verbs). The X-axis represents time from –200 ms (before stimulus onset) to 2000 ms, with each tick mark representing 250 ms. The Y-axis shows voltage from –2 μV to 2 μV, with negativity plotted downward.

### Korean Active Group: Semantically Violated **–** Control

The N400-like is observed in in the centro-parietal and occipital region between approximately 350 and 650 ms. Interestingly, positivity started from 350 ms and peaked droung 750 ms was also observed in right, left and tempro parietal and occipital regions (Fig. 4B).

### Chinese Sham Group: Semantically Violated **–** Control

In topographic Fig. 4C, the difference wave is calculated by subtracting the ERPs elicited by the control condition from those derived from the semantic condition in sham Chinese participants (semantic – control). An N400-compatible negativity appeared in fronto-central sites (350–449 ms), followed by a broad P600-compatible positivity over centro-parietal areas between 450 and 650 ms.

### Chinese Active Group: Semantically Violated **–** Control

The N400 is more focused and pronounced in the centro-parieto-occipital regions between approximately 251 and 549 ms. Also, a positivity was observed in the left fronto-parieto-temporal regions that was extended to the right fronto-parieto temporal around 651-750 ms (Fig. 4D).

### N400 (350–550 ms)

A linear mixed-effects (LME) model (fit by REML) was used to analyze N400 amplitudes. No significant effect of stimulation type (sham – active: *β* = –2.68 × 10⁻⁷, *p* = .21) nor L1 (Korean – Chinese: *β* = 6.98 × 10⁻⁷, *p* = .15) was found. However, session number had a significant positive effect on amplitude (*β* = 2.20 × 10⁻⁷ per session, *SE* = 6.20 × 10⁻⁸, *t* (960) = 3.54, *p* < .001). The model also revealed a significant L1 × sentence type’s interaction (*F* (5, 960) = 2.28, *p* = 0.045), a stimulation × L1 interaction (F (1,960.7) =5.48, p=.0194), as well as stimulation × sentence type (*F* (5,960.7) = 2.28, *p* = .0448).

Compared to Chinese participants, Korean participants exhibited reduced N400 amplitudes for semantic violated sentences (transitive verb type) (*β* = −8.374e−7, *p* = 0.005) and semantic violated sentences (intransitive verb type) (*β* = −6.107e−7, *p* = 0.036).

### P600 (600–900 ms)

Conversely, there was no significant effect of session number (*F* (1,1302.7) = 1.16, *p* = 0.28), stimulation (sham – active: *β* = 1.47 × 10⁻⁷, *SE*=1.54 × 10⁻⁷, *t*(1303) = 0.96, *p* = 0.339), L1 (Korean – Chinese: *β* = 3.52 × 10⁻⁷, *SE* = 5.53 × 10⁻⁷, *t*(28)=0.64, *p* = 0.529), nor any single sentence type contrast was significant. However, a significant effect of stimulation × L1 interaction was shown (*F* (1,1303.1) = 9.86, *p* = 0.0017), stimulation × sentence type (*F* (5,1303.1) = 2.35, *p* = 0.039), and L1 × sentence type’s interaction (*F* (5,1303.1) = 4.53, *p* < 0.001).

### Grand Average ERPs in the ROI Centro-parietal region

## Discussion

The present tDCS-EEG study explored the neural mechanisms involved in processing syntactic and semantic anomalies in L2 among Korean and Chinese native speakers fluent in Japanese. Given that Japanese and Korean share morpho-syntactic features, and Japanese and Chinese share Chinese characters with semantic radicals, we aimed to investigate the L1 transfer effects.

The analysis of accuracy revealed significant effects of tDCS stimulation of the LIFG and LSTG and L1 transfer impact on processing performance. For Chinese speakers, semantic violations led to significantly higher accuracy than control and syntactic violations. This indicates an L1 Chinese Kanji (which has semantic radicals) effect transferring to L2 Japanese learners. Interestingly, semantic violations also led to significantly higher accuracy for Korean speakers than syntactic violations and control conditions; however, there was no significant difference between control and semantic conditions. This might suggest that the increased difficulty of syntactic violations, combined with the presence of Japanese Kanji semantic radicals, could lead to a higher cognitive load for Korean speakers, resulting in lower accuracy given the short time required to answer.

For RT, intransitive verbs showed larger RT differences than transitive verbs in the significant condition × transitivity interaction. In particular, syntactic processing – semantic processing (Δ*M* = 0.3292 for intransitives vs. 0.1825 for transitives) and control processing – semantic processing (Δ*M* = 0.2248 for intransitives vs. 0.0590 for transitives) differed. Whereas Korean participants showed no significant transitivity-related RT differences (*p* > 0.05), Chinese participants exhibited significant transitivity effects, with slower RTs for intransitive verbs compared to transitive verbs in both control (*p* = 0.005) and syntactic conditions (*p* = 0.007). Moreover, semantic violations generally reduced transitivity effects across both groups, emphasizing the relative efficiency of semantic over syntactic integration in L2 contexts. These behavioral results underline significant cross-linguistic differences: Chinese participants struggle more with intransitive constructions that lack supportive morphological cues, whereas the L1’s case-marking system in Korean influences how they efficiently process both verb types— transitive and intransitive.

Using dual active tDCS stimulation over the LIFG and LSTG combined with EEG, the findings from ERPs revealed L1-specific modulations in the L2 P600 and N400 components associated with semantic integration and syntactic reanalysis, respectively. Dual active stimulation led not only to amplified ERP markers but also narrowed their distribution to task-related regions and time windows.

For syntactic violations, active dual-site tDCS produced a more sustained P600 over centro-parietal regions (450–950 ms for Koreans; 450–849 ms for Chinese), coupled with attenuated early N400-compatible negativities, suggesting facilitated syntactic reanalysis relative to sham (Figs. 6, 8, 10, 12). The observed N400-compatible negativity in the centro-parietal region for Korean and Chinese participants under sham and active stimulation in the syntactically violated sentences, peaking around 251–449 ms, suggests an initial difficulty in interpreting syntactic information.

**Fig. 6.**
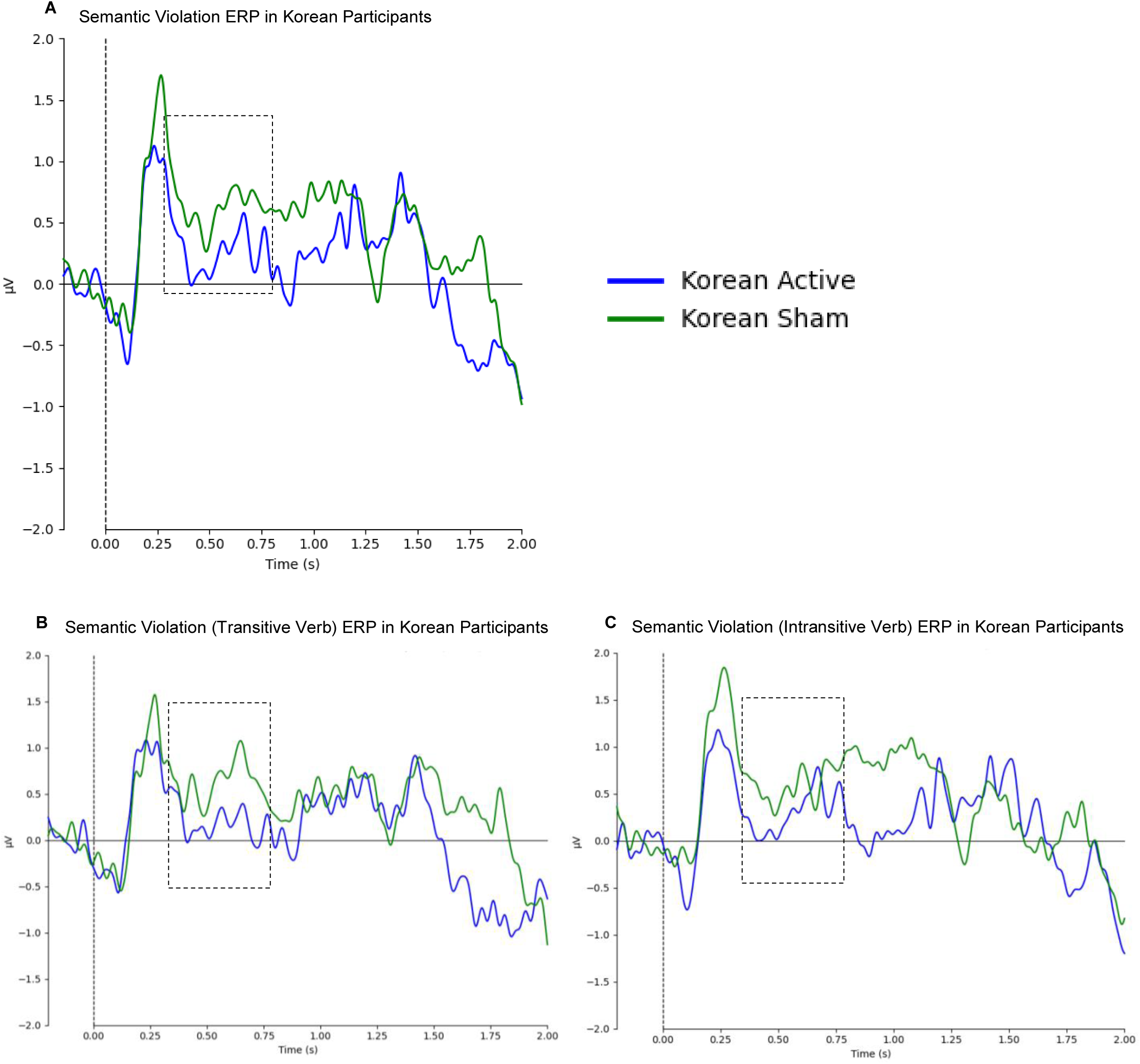
Mean grand average ERPs in Centro-parietal region in semantic violated sentences in Korean participants (n = 12). **A**, Sham stimulation (green) shows a positivity (P600) around 500–700 ms, while under active stimulation, an N400-like waveform is observed around 300–500 ms. **B**, The P600 is significantly reduced, and the N400 is increased in the active stimulation group (transitive verbs). **C**, Active stimulation reduces the P600 and increases the N400 in the intransitive verbs. The X-axis represents time from –200 ms (before stimulus onset) to 2000 ms, with each tick mark representing 250 ms. The Y-axis shows voltage from –2 μV to 2 μV, with negativity plotted downward.

**Fig. 7.**
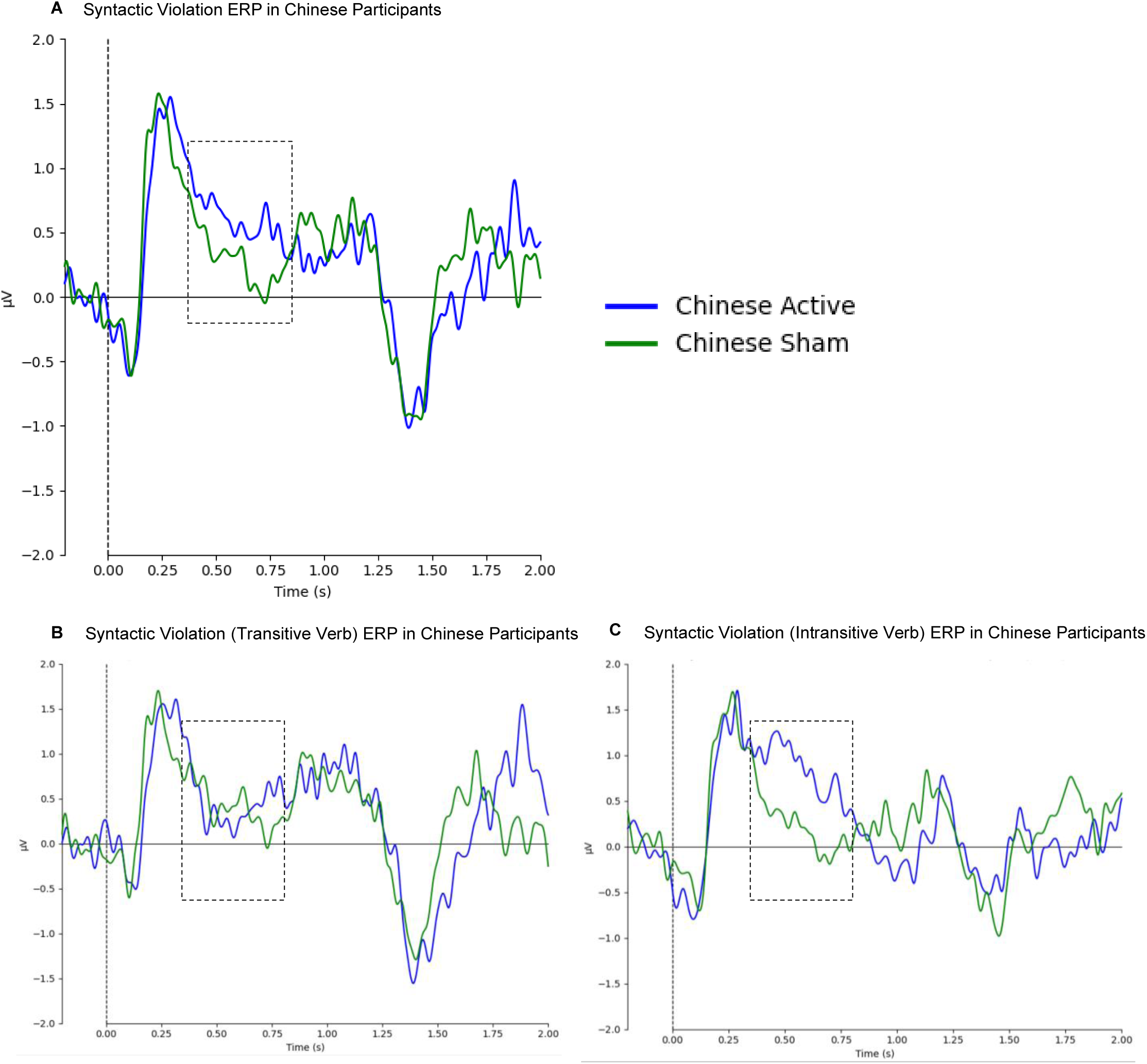
Mean grand average ERPs in Centro-parietal region in syntactic violated sentences in Chinese participants (n = 15). **A**, Active stimulation (blue) shows a P600 around 500–700 ms and reduces the negativity observed around 300–500 ms in the sham group. **B**, Active stimulation shows a negativity (N400) around 300–500 ms and a reduced P600 during processing of syntactically violated sentences (transitive verbs). **C**, Active stimulation shows a reversed positivity of the negative waveforms observed around 300–700 ms in the sham group (intransitive verbs). The X-axis represents time from –200 ms (before stimulus onset) to 2000 ms, with each tick mark representing 250 ms. The Y-axis shows voltage from –2 μV to 2 μV, with negativity plotted downward.

**Fig. 8.**
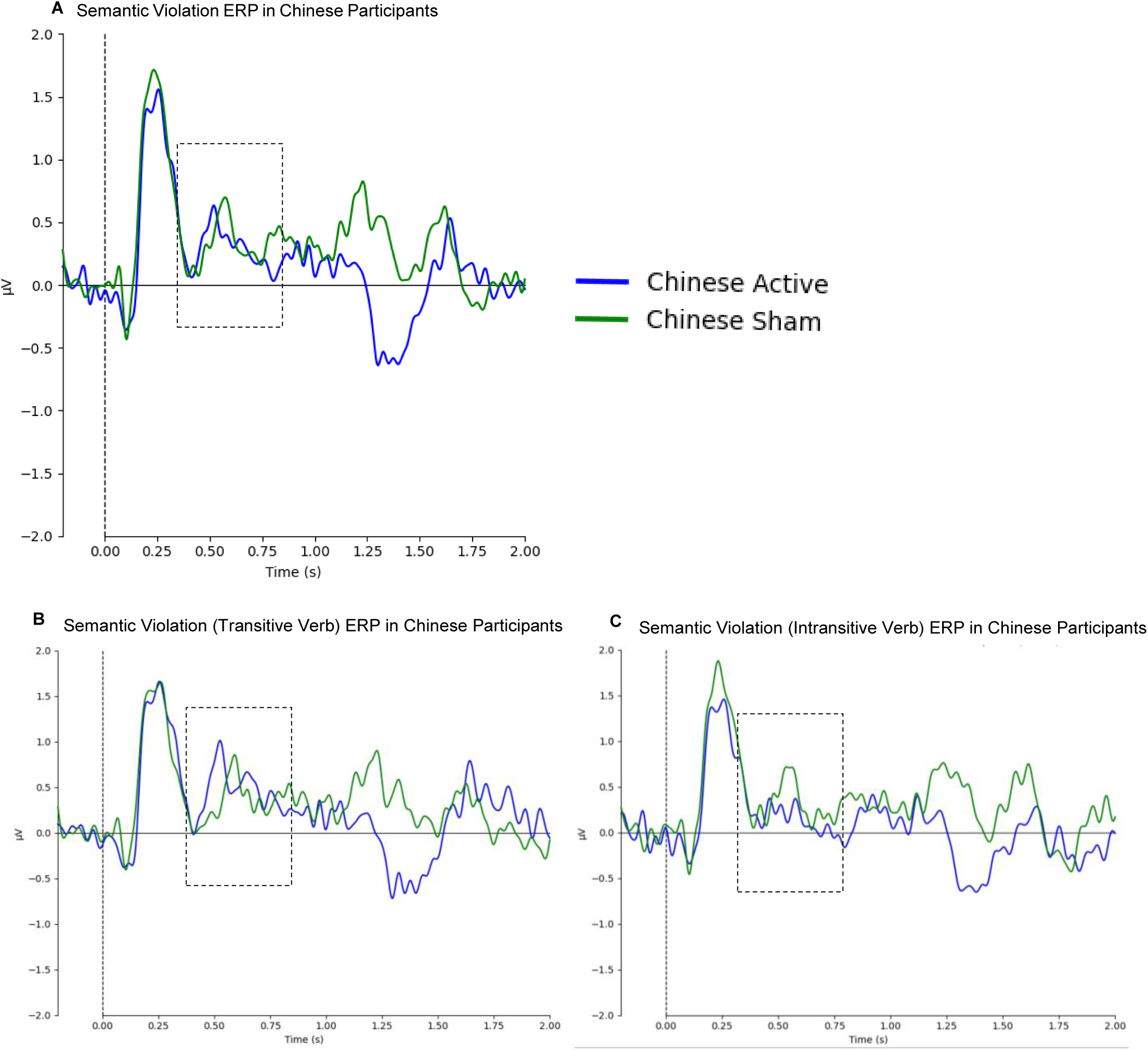
Mean grand average ERPs in Centro-parietal region in semantic violated sentences in Chinese participants (n = 15). **A**, Under sham stimulation (green), a slight N400-like waveform is observed around 300–500 ms, which is reduced in the active stimulation group, showing a P600 peak around 550–700 ms. **B**, The P600 peaks slightly earlier under active stimulation, while the N400 is similar in both conditions (transitive verbs). **C**, Active stimulation reduces the P600 (intransitive verbs). The X-axis represents time from –200 ms (before stimulus onset) to 2000 ms, with each tick mark representing 250 ms. The Y-axis shows voltage from –2 μV to 2 μV, with negativity plotted downward.

This may mean that L2 speakers experience challenges retrieving word meanings, integrating them into context, and processing sentence meaning—especially when encountering unfamiliar or complex vocabulary. The P600 was observed in the left centro-temporo-parietal regions, suggesting that both the sham and active stimulation groups engaged in cognitive processes related to syntactic repair or reanalysis.

For semantic violations, active stimulation yielded a somewhat larger N400 in centro-parietal sites (350–750 ms for Koreans; 251–549 ms for Chinese), along with a reduced P600, consistent with enhanced semantic integration under dual-site tDCS (Figs. 7, 9, 11, 13). The LME model showed that N400 dynamics—indicating accumulated training (N400 increasing with session number)—enhanced semantic integration, and P600 amplitudes indicated L1-modulated syntactic processing. However, the relatively small sample size may limit the statistical power to detect subtle interaction effects and may reduce the generalizability of the findings across broader L2 populations.

**Fig. 9.**
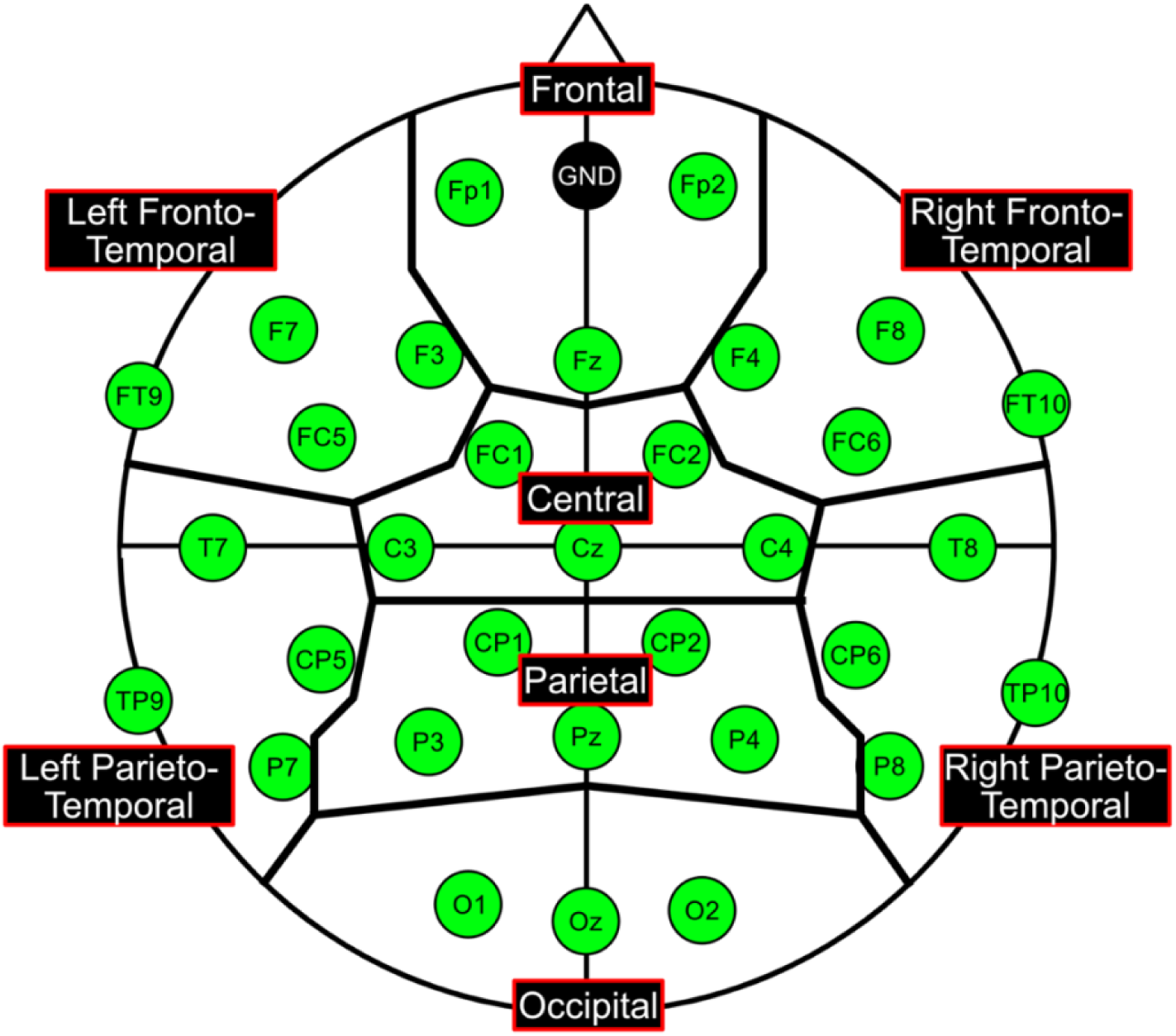
ROIs for the N400 and P600. The spatio-temporal analysis of the P600, we used electrodes within the parietal area as the regions of interest (ROIs), whereas for the N400, we used electrodes within the centro-parietal area as the ROI.

To sum up, the results suggest that active tDCS stimulation modulates ERP magnitudes and shifts the topography to more specific, limited regions and time windows compared to sham stimulation. This effect is particularly evident in the enhanced P600 component, which illustrates improved cognitive functioning and engagement in processing both syntactic and semantic violations. Dual active tDCS stimulation of the LIFG and LSTG can modulate brain activity and improve language comprehension in L2 speakers. Moreover, the behavioral data show that semantic anomalies were consistently linked to higher accuracy among Chinese and Korean participants. This may imply that semantic processing is more straightforward for L2 speakers than syntactic processing.

Future research is encouraged to explore the effects of various types of stimulation, such as transcranial alternating current stimulation, and their long-term impacts on language learning and cognitive processing in both L1 and L2 speakers. Additionally, investigating other language groups and more complex linguistic structures and sentences could provide further insights into the generalizability of these findings.

## Materials and Methods

### Participants

We recruited 15 Chinese (5 males) aged 21–33 years (mean ± SD = 24.93 ± 3.86) and 12 Koreans (9 males) aged 20–33 years (mean ± SD = 25.92 ± 4.52), all native-speaker undergraduate and graduate students fluent in Japanese at Kyushu University in Fukuoka, Japan. All participants held JLPT N1 certification except one Chinese student, who held JLPT N2 certification; native Japanese speakers were excluded. Only right-handed participants were included; right-handedness was confirmed using the Edinburgh Handedness Inventory (Oldfield, 1971; mean laterality quotient = 86.67). Participants had no history of neurological, speech, or mental disabilities. Normal or corrected-to-normal visual acuity and hearing were required, allowing the use of glasses or contact lenses. Informed consent was obtained from each participant after a full explanation of the experiments, in accordance with the World Medical Association’s Declaration of Helsinki. The experimental procedures were approved by the Ethics Committee of the Faculty of Humanities at Kyushu University.

The overall experiment duration was approximately three hours, with data collection expected to be completed within one hour. Each participant took part twice, with approximately a three-week interval between sessions. Sessions were randomly assigned as dual-site anodal or sham tDCS, ensuring that each participant experienced both conditions (sham and active). We also ensured that the number of first-session sham receivers was similar to the number of first-session active receivers. A single-blind, sham-controlled, within-group design was used—that is, participants were blinded to the randomization, while the experimenter was not.

### Stimuli

Stimuli were created by a native Japanese linguist and revised by two other native Japanese linguists. Stimuli were further checked by a non-native Japanese speaker fluent in Japanese to verify the difficulty level and double-check sentence design from the perspective of second-language Japanese learners. A total of 198 target sentences (3-word sentences consisting of two noun phrases and a verb) and 66 filler sentences were presented to prevent participants from guessing the purpose of the experiment and to ensure unbiased responses. The sentences were categorized into three conditions: correct, syntactically violated, and semantically violated. All experimental stimuli (including both targets and fillers) are presented in the Appendix.

### Correct sentences

First, we used 66 control correct sentences, divided into 33 with transitive verbs and 33 with intransitive verbs.

|  |  |  |
| --- | --- | --- |
| 子供が | 虫を | 見つける |
| Kodomo ga | musi o | mitukeru |
| Child NOM | insect ACC | findtransitive |
| “A child finds an insect.” |  |  |

### Syntactically violated sentences

Second, we made 66 syntactically violated sentences, divided into 33 with transitive verbs and 33 with intransitive verbs. To make these syntactically violated sentences, we shuffled the verbs from the correct sentences, transitive verbs to intransitive verbs and vice versa, while keeping the argument structure and the case markers. For example, the above control sentence “kodomo ga musi o mitukeru” was changed as follows.

|  |  |  |
| --- | --- | --- |
| *子供が | 虫を | 見つかる |
| Kodomo ga | musi o | mitukaru |
| Child NOM | insect ACC | findintransitive |

### Semantically violated sentences

Third, we prepared 66 semantically violated sentences, divided into 33 with transitive verbs and 33 with intransitive verbs. We shuffled the verbs in the correct sentences within transitive or intransitive verbs to violate the meaning of the sentence. We did not shuffle intransitive verbs to transitive verbs or vice versa to prevent a syntactic violation in the semantically violated sentences.

|  |  |  |
| --- | --- | --- |
| #子供が | 虫を | 閉める |
| Kodomo ga | musi o | simeru |
| Child NOM | insect ACC | close |

### Procedure

Sentence correctness or naturalness judgment tasks were conducted using PsychoPy (version 3.0.7), a Python-based psychophysics software library (Peirce, 2007). We used the Meiryo font, colored in grey, to reduce eye discomfort caused by the high contrast of black and white. Each trial started with a blank black screen followed by a fixation cross (“+”) for 500 milliseconds (ms), a first noun phrase (NP) for 1000 ms, a second NP for 1000 ms, and a verb for 1000 ms (Fig. 1). After the verb disappeared, a task screen was displayed for up to 4000 ms or until the participants answered. The task was to judge the naturalness of the sentences by pressing the right button if the sentence was natural and the left button if it was unnatural.

A total of 263 sentences were randomly shown in nine sessions, with a short break between each session of about 30 sentences. Participants could press “continue” whenever they were ready to proceed. The total average time of the short breaks for all participants was approximately one minute. The trial order was randomized for each participant.

Participants sat in a comfortable chair in a dimly lit, electromagnetically shielded room. A liquid crystal monitor was positioned about 130 cm in front of them, with the chair and screen aligned as precisely as possible. While placing the EEG electrodes on the cap, along with the necessary dual-site tDCS electrodes, participants were offered the option to watch a movie or YouTube, or simply rest, to ensure they were relaxed and not bored before the experiment started. After completing all electrode installations, tDCS stimulation began immediately and lasted for 20 minutes. Participants were instructed not to sleep or make sudden or aggressive head movements. To ensure their physical condition remained stable and that they followed instructions properly, they were monitored throughout the entire experiment using an insight-monitoring camera.

Once the tDCS stimulation was over, we checked participants’ readiness to start the main experiment. The PsychoPy program was designed to show five practice trials that included all three conditions (control, syntactic violation, and semantic violation). Feedback on the practice trial answers was given to participants to ensure they could judge the sentences correctly and understand the pacing of phrase presentation.

### EEG and tDCS

First, we marked the Cz electrode position following the international 10-20 system. Then, the LIFG (corresponding to electrode FC5) and LSTG (corresponding to electrode TP7) regions were marked with a skin marker.

Because this was a 2 × 2 stimulation design, we used two types of electrodes. The first set, which served as cathodes, consisted of two large rubber-ring electrodes (outer diameter 75 mm, inner diameter 30 mm, thickness ∼2 mm). They were placed over the marked LIFG and LSTG regions, allowing the markings to remain visible through the inner holes of the electrodes. Without moving the electrodes, the EEG cap was carefully placed on the participant’s head.

Next, we used three Ag/AgCl electrodes (outer diameter 12 mm, inner diameter 5 mm, thickness 2 mm). Two of these were placed at FC5 and TP7, over the previously marked sites. The third electrode was placed at CPz and used as a reference. HD-tDCS stimulation was conducted using the Soterix Medical HD-tDCS system (Soterix Medical Inc., Woodbridge, NJ, USA). The protocol was configured as follows: channel 1 set to +1 mA (anode), channel 2 set to –1 mA (cathode), channel 3 set to +1 mA (anode), and channel 4 set to –1 mA (cathode), resulting in a total of 2 mA of anodal stimulation. For the anodal tDCS group, 2 mA was applied simultaneously for 20 minutes, including 30-second ramp-up and ramp-down periods to minimize skin sensation over the LIFG and LSTG (Fig. 2B).

The EEG data were recorded from 32 active electrodes (actiCAP, Brain Products) and amplified on a Neurofax EEG-1200 system (Nihon Kohden). Four additional Ag/AgCl electrodes were placed as follows: two were positioned below and to the left of the left eye to track vertical and horizontal eye movements; two were connected to the earlobes (A1 and A2) and served as reference electrodes; and one was placed at Fpz to provide a baseline signal for stabilizing the EEG recording. The impedance of all electrodes was maintained below 45 kΩ throughout the experiment. EEG was sampled at 1,000 Hz after being bandpass filtered from 0.78 to 120 Hz.

## Data Analyses

### Behavioral Data

For behavioral data of accuracy and reaction time were analyzed using R Studio (2024.09.0) and the function of “anovakun” (version 4.8.9), with a fixed 4-second interval between the final phrase (verb presentation) and participant’s response. We ran a four-way mixed-design analysis of variance (ANOVA) (Stimulation type [active, sham] x Sentence condition type [control, semantic violation, syntactic violation] x Language [Chinese, Korean]) x Verb type [transitive and intransitive] (significance level α = 0.05). The Stimulation, Sentence condition type and Verb type are within-subject factors, while Language is a between-subject factor. If the main effects or interaction were significant, post-hoc comparisons using Shaffer’s Modified Sequentially Rejective Bonferroni (Shaffer’s MSRB). Procedure was also performed to evaluate pairwise differences. If the sphericity assumption was violated, the Greenhouse-Geisser correction was also applied for the within-subject factors to adjust the degrees of freedom. In addition, the aggregated data files were exported to JASP (Version 0.19.1) for plotting graphs. Trials with missing and incorrect responses were excluded from Anova analyses. In addition, trials with standard deviation 3 standard deviations above the mean of reaction times (RT) were excluded.

### EEG Data

EEG data was processed using MNE-Python 1.4.0 along with custom Python scripts. All analyses were performed in Python 3.12.2 (64-bit) using the Spyder IDE (Anaconda distribution), and the processing steps were as follows. First, event lists were created to identify the trigger events in the EEG data, providing an indicator of when each trigger occurred. Second, bad channels were manually selected, removed, and interpolated. Third, EEG data was automatically preprocessed by filtering and referencing using a high-pass filter at 0.1 Hz, and eye blink artifacts were removed using independent component analysis (ICA). Subsequently, a step involving epoching, rejecting noisy trials, and saving logs of cleaned data for each participant was performed. Fourth, by applying a low-pass filter at 30 Hz, we conducted a grand averaging of ERPs across all trials for all conditions and participants.

For ERP analysis, we obtained N400 and P600 components. The mean N400 and P600 amplitude within the window of interest (N400: 350∼550 ms, P600: 600∼900 ms), per ROI (Fig. 9) and participant was calculated.

### Statistical analysis

ERP data for the N400 and P600 components were analyzed using LME models in R studio. Mean amplitude values were extracted on a trial-by-trial basis. For the N400 analysis, data were filtered to include the data in the centroparietal region electrodes only. For the P600, only the parietal region electrodes were included. Mixed-effects models were fitted using the lmer function from the lme4 package, where the fixed effects were stimulation (sham or active), L1 (Korean or Chinese), sentence type (correct, semantic violated or syntactic violated), session (first or second), and the random intercept was Participant [Mean Amplitude ∼ Stimulation * L1 * Condition + Session + (1 | Participant)]. The guidance for model selection was done by stepwise comparison and Akaike’s Information Criterion (AIC) values. Post-hoc pairwise comparisons were also conducted using estimated marginal means (EMMs) with Bonferroni correction.

## Notes

### Competing Interest Statement

The authors have declared no competing interest.

